# State- and Identity-Dependent Motor Neuron Excitability Shapes Cutaneous Long-Latency Reflexes

**DOI:** 10.64898/2026.03.25.714138

**Authors:** Yannick Finck, Demetris S. Soteropoulos, Alessandro Del Vecchio

## Abstract

Neuromuscular reflexes elicited by sensory nerve stimulation provide valuable insights into neural motor control pathways. Analysis at the level of individual motor units (MUs) is feasible via electromyographic decomposition, but the factors shaping MU-specific reflex responses remain poorly understood. We investigated long-latency responses to cutaneous electrical stimulation in a large population of tibialis anterior MUs from nine healthy subjects during isometric ankle dorsiflexion at 10–30% of maximum voluntary contraction. Individual MU reflex responses differed markedly. Using 1000 stimulation pulses per trial, substantially more than the 150–300 typically reported in previous studies, provided more reliable estimates of cutaneous reflex characteristics. Across the motor pool, reflex magnitude increased with force level (*p* < 0.001) while excitation probability correlated significantly with MU recruitment threshold in 78% of subjects (*p* = 0.012). Furthermore, excitation probability increased systematically with contraction intensity (*p* < 0.001) for individually tracked MUs. Post-excitatory depression (PED) magnitude correlated significantly with excitation probability (*r* = 0.50, *p* < 0.001) of individual MUs. A targeted reflex-removal analysis, validated by MU simulations incorporating realistic excitation probabilities into ordinary firing patterns, reduced the PED by 84.2% in simulated data but only by 34.7% in recorded units. These findings suggest that the PED is a complex, hybrid phenomenon, resulting from synchronization-induced discharge resetting and additional independent inhibitory components. These findings demonstrate that MU-level reflex excitability to somatosensory input is influenced by state- and identity-dependent motor neuron characteristics, underscoring the importance of using sufficient stimulation pulses for reliable reflex measures and MU population analysis.

## I. Introduction

The analysis of reflex responses is a fundamental approach for probing human neuromuscular pathways and understanding the integration of sensory input into motor output [1]. Central to this process are long-latency reflexes (LLRs), involuntary muscle activations that occur after the spinal short-latency reflex (SLR) but before the onset of voluntary movement [2]. These responses are widely regarded as a sophisticated mechanism of sensorimotor integration [3], reflecting the nervous system’s ability to modulate motor output based on task demands and sensory feedback [4].

Historically, the scientific understanding of LLRs evolved from the study of the stretch reflex. While early 20th-century research focused on the spinal “tonic” components of reflex activity [5, 6], subsequent investigations identified that the reflex response to muscle stretch comprises multiple components: an early spinal M1 response and longer-latency M2 and M3 components [4, 7, 8]. The hypothesis that these longer-latency components involve a transcortical loop, traversing the sensorimotor cortex, was later confirmed through landmark studies demonstrating that the integrity of the dorsal columns and the motor cortex is essential for their expression [9–12].

Critically, these transcortical circuits are not only engaged by proprioceptive input but are also highly responsive to cutaneous afferents [13–16] Systematic work by Deuschl and colleagues further unified these observations, characterizing LLRs as involuntary reactions that follow a similar temporal pattern regardless of whether the triggering stimulus is a muscle stretch, electrical nerve stimulation, or a complex cutaneous input [2]. Consequently, the analysis of these reflexes has become a cornerstone in clinical neurophysiology, serving as a valuable tool for supporting diagnosis, monitoring disease progression, and probing the underlying neural circuits of sensorimotor control [17].

Against this background of historical and clinical interest in LLRs, little is known how populations of spinal motor neurons from an individual motor nucleus integrate processing of the cutaneous LLRs across different healthy human subjects during different voluntary muscle activation levels. In the present work, we investigated the responses of different populations of spinal motor neurons in the tibialis anterior muscle to cutaneously-elicited LLRs. Specifically, we investigated the reflex response following electrical stimulation of cutaneous afferents innervated by the fibular nerve (superficial or common) at the dorsum of the foot. The shape of the average cutaneomuscular reflex response in the EMG signal of the tibialis anterior is similar to previously reported responses [16, 20, 21], which typically include a strong long-latency component in the form of excitation at approximately 70-100 ms post-stimulus, followed by an oscillatory post-excitatory wave (PEW) that gradually decays towards baseline (Figure 1b).

**Fig. 1:**
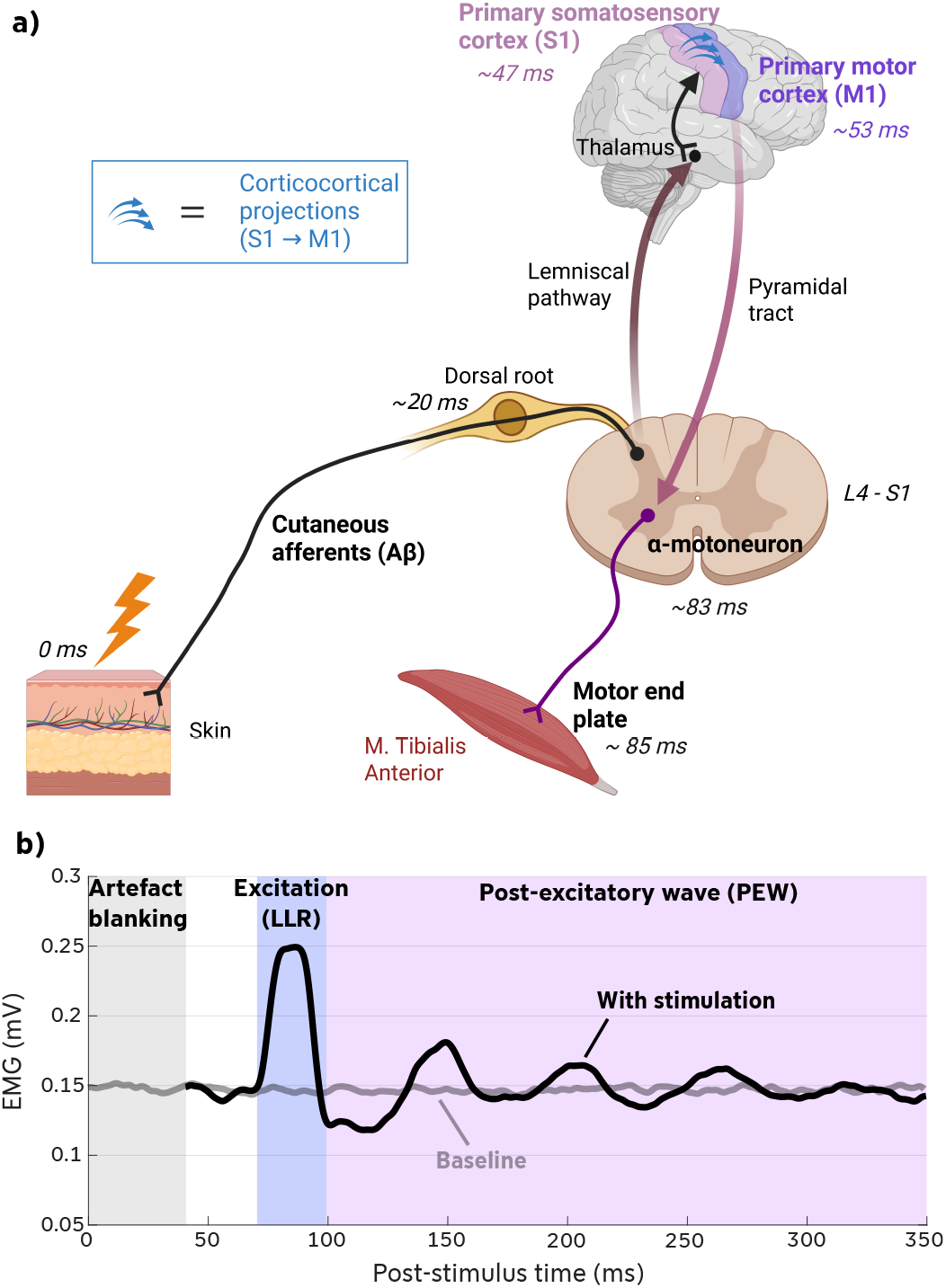
a) Transcortical pathway of a LLR evoked by cutaneous stimulation of sensory nerves. According to some definitions, LLRs also include spinal loops [17] which were omitted in this illustration to emphasize the transcortical pathway. The approximate conduction and processing times indicated at each station are based on data from [18, 19]. b) Exemplary cutaneomuscular reflex. The dark black trace shows the typical shape of the post-stimulus averaged electromyography (EMG) response of the tibialis anterior muscle following cutaneous electrical stimulation of the superficial fibular (peroneal) nerve during a constant voluntary contraction. Between approximately 70 and 100 ms, the response is characterized by the excitation associated with the LLR, followed by a oscillatory post-excitatory wave (PEW). The light grey trace represents the baseline EMG recorded without stimulation. The first portion of the stimulated EMG signal (up to ∼50 ms) is usually omitted due to stimulation artefacts.

Previous studies of these cutaneously-mediated LLRs have been predominantly limited to the analysis remaining at the EMG-level [13, 22–2 While global EMG metrics provide an aggregated measure of muscle activity, they are confounded by amplitude cancellation and fail to reveal the dynamic responses of individual motor units (MUs) [25, 26]. Therefore, signal decomposition is necessary to analyze the neural drive accurately. Understanding how individual MUs react to the sensory input is crucial for fully comprehending the central mechanisms of sensorimotor integration and the interpretation of excitatory and inhibitory pathways [27–29].

Our objective is therefore to characterize the reflex responses to a cutaneous stimulus at the level of single MUs. We investigate whether MUs exhibit different reflex responses and which factors and MU-specific characteristics may influence these responses. Previous studies that have conducted similar MU-level analyses have, in our estimation, often employed an insufficient number of stimuli for reliable estimation of reflex parameters. The typical number of stimuli used to calculate LLR parameters of single MUs in these works was in the range of approximately 150 to 300 [1, 16, 21, 30, 31]. This range aligns with figures typically reported in the literature; for instance, a comprehensive review on LLRs reported stimulus counts often found to be between 64 and 256 repetitions for reliable analysis [17]. Therefore, an additional aspect of this work is the methodological examination of the stability and reliability of these MU-specific reflex parameters as a function of a significantly increased number of stimuli. We hypothesize that a substantially increased number of repetitions is required to reduce variability and enable robust estimation of reflex response parameters at the level of individual motor neurons. Our primary hypothesis is that LLR responses are shaped by the combined integration of spinal motor neuron excitability and the baseline net neural drive, which together determine the expression of cutaneous reflex responses. We further hypothesize that LLR modulation at the level of individual motor neurons follows a continuum consistent with Henneman’s size principle, such that smaller and larger motor neurons exhibit systematically graded responses. The analysis of motoneuron populations then allowed us to refine new a posteriori hypothesis, leading to a clearer understanding of the PEWs.

In summary, this study aims to advance the understanding of cutaneously-elicited LLRs in three critical ways. Firstly, we provide a high-resolution analysis of LLR modulation by characterizing the response heterogeneity of individual MUs in the tibialis anterior across different levels of voluntary background contraction and determining which MU-specific characteristics (e.g., recruitment threshold or firing properties) correlate with this variability. Secondly, we address a significant methodological limitation in the field by systematically investigating the impact of an increased number of stimuli on the stability and reliability of MU-specific reflex parameters. We hypothesize that a substantial increase in stimulus count is required to achieve robust estimates. Thirdly, we investigate the underlying mechanism of the PEW, which typically follows the LLR excitation in the MU firing pattern. Specifically, we test the hypothesis that the entire subsequent declining PEW visible in the EMG signal and post-stimulus time histogram (PSTH) is primarily caused by a reset and therefore synchronization of the MU firings due to a synchronization-induced post-excitatory depression (PED) rather than being generated by multiple, distinct components of a centrally mediated reflex pathway. We use the area of the initial PED as an indicator for the magnitude of the PEW.

## II. Materials and Methods

### A. Participants

Nine healthy volunteers, including two females and seven male participants, aged 24 to 28 (referred to as S1-S9), participated in the study. For all subjects, EMG signals were acquired using high-density surface electrodes (high-density surface electromyography (HDsEMG)). The study protocol was identical for all participants. The study was approved by the ethics committee of the Friedrich-Alexander-University Erlangen-Nürnberg (approval number 25-37-S) and complied with the Declaration of Helsinki. All participants provided written informed consent prior to their participation.

### B. Electrical Stimulation

Electrical stimulation was delivered using a DS8R biphasic constant current stimulator (Digitimer, UK) connected to ValuTrode cloth electrodes (Axelgaard, USA), with a round cathode (2.5 cm diameter) and an oval anode (4x6 cm). The electrodes were affixed to the skin of the right dorsal foot. The cathode was positioned precisely over the bifurcation point where the superficial fibular (peroneal) nerve, specifically the medial dorsal cutaneous branch, divides into the dorsal metatarsal and digital nerve branches. The anode was placed in the flexion crease between the foot and the lower leg (i.e., the anterior ankle region).

Stimulation was applied with a pulse width of 100 *µ*s and a current amplitude of approximately 2.5 times the individual perceptual threshold of each subject [22]. Stimuli were delivered as single pulse stimuli at a frequency of approximately 2 Hz, with inter-stimulus intervals uniformly distributed between 350 ms and 650 ms [32] (Figure 2b) to prevent adaptation to the stimulus.

**Fig. 2:**
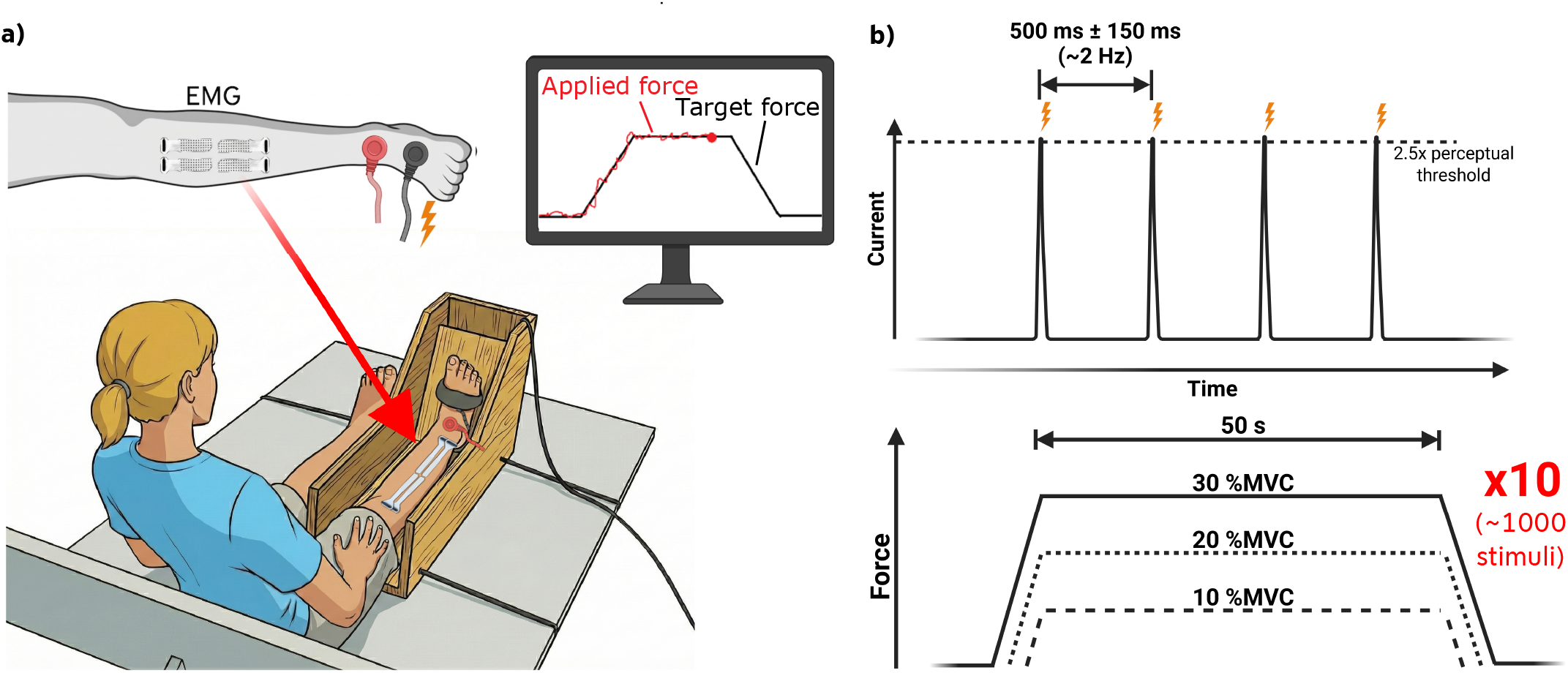
a) Experimental setup. Four HDsEMG electrode grids were placed over the Tibialis Anterior muscle, and stimulation electrodes were attached to the dorsum of the foot. Participants sat upright with extended legs, and the right foot was secured in the ankle dorsiflexion dynamometer. Real-time force feedback was provided to the participant via a monitor. b) Stimulation parameters (top) showing electrical stimulation at approximately 2 Hz and 2.5 times the individual perceptual threshold regarding the current level. Experimental protocol (bottom) with trapezoidal force trajectories at 10, 20, and 30 % of maximum voluntary contraction (MVC). Each target force level was repeated 10 times with stimulation enabled during the ramp execution phase.

### C. Data Acquisition

HDsEMG signals were recorded from the tibialis anterior muscle of the right leg. For HDsEMG, four 13×5 electrode grids (each comprising 64 electrodes, totaling 256 electrodes) with an interelectrode distance (IED) of 4 mm were placed on the skin over the muscle belly of the tibialis anterior in a 2×2 matrix aligned longitudinally with the muscle fibers (Figure 2a). A conductive paste was applied between the grids and the skin to improve electrode-skin contact. surface electromyography (sEMG) signals were sampled at 2048 Hz and were amplified and acquired using a Quattrocento amplifier system (OT Bioelettronica, Italy).

To record dorsiflexion force, a custom-built ankle dorsiflexion dynamometer (OT Bioelettronica, Italy) was used to measure isometric force output during right ankle dorsiflexion.

### D. Experimental Protocol

Participants were seated in an upright position on an examination table, with both legs extended in a relaxed and comfortable posture. The right leg was positioned in the ankle dorsiflexion dynamometer, and the foot was securely fixated to the force sensor just distal to the ankle joint (Figure 2a). To assess MVC, participants were instructed to perform three maximal isometric ankle dorsiflexions, and the highest value was used for normalization.

Stimulation electrodes were then applied to the skin as previously described. Electrical stimulation was first tested to allow participants to familiarize themselves with the sensation. The individual perceptual threshold was determined by gradually increasing the stimulation current until the first sensation was perceived, and subsequently decreasing it until the perception disappeared completely. For safety, it was verified that the stimulation amplitude used during the recordings, 2.5 times the perceptual threshold, remained below the pain threshold for each participant. This condition was met in all cases. After the stimulation setup, the EMG electrode grids were adhered to the skin over the tibialis anterior as previously described.

Following preparation, the main experimental protocol commenced. During each trial, participants were instructed to follow a trapezoidal force trajectory displayed on a screen by dorsiflexing their right ankle. Each trapezoidal ramp consisted of a 50-second plateau phase with linear ascending and descending phases of 5 seconds each. To achieve an average of approximately 1000 delivered electrical stimuli during the plateau phases, a total of 10 ramps were performed per recording session, with 30-second rest periods between ramps (Figure 2b) to prevent muscle fatigue.

A familiarization trial consisting of a single ramp was conducted first. Subsequently, three recording sessions were performed at plateau levels of 10, 20, and 30 %MVC. Electrical stimulation was delivered during each ramp as previously described and was deactivated during the rest phases. The three contraction levels were performed in random order. A 5-minute break was provided between recording sessions to prevent muscle fatigue.

### E. Data Analysis

For each recording, individual MUs were identified from the EMG signals using a blind source separation (BSS) technique [33, 34] that provides fully automatic decomposition. Prior to decomposition, the EMG signals were digitally band-pass filtered to reduce noise and preserve the physiological frequency content. For HDsEMG, a band-pass filter with cut-off frequencies of 20–450 Hz was applied, downsampled at 2048 Hz. MUs were only included in the analysis if their decomposition quality exceeded a silhouette value (SIL) of 0.9 [34] and undercut a coefficient of variation (CoV) of inter-spike interval (ISI) values of 0.5. Each recorded file, comprising all 10 ramp contractions, was decomposed as a whole.

The intensity of the elicited transcortical reflex was quantified using a PSTH. In this study, the PSTH reflects the probability of MU discharges within specific time intervals following an electrical stimulus (Figure 3). Two different computation approaches were used:

**Fig. 3:**
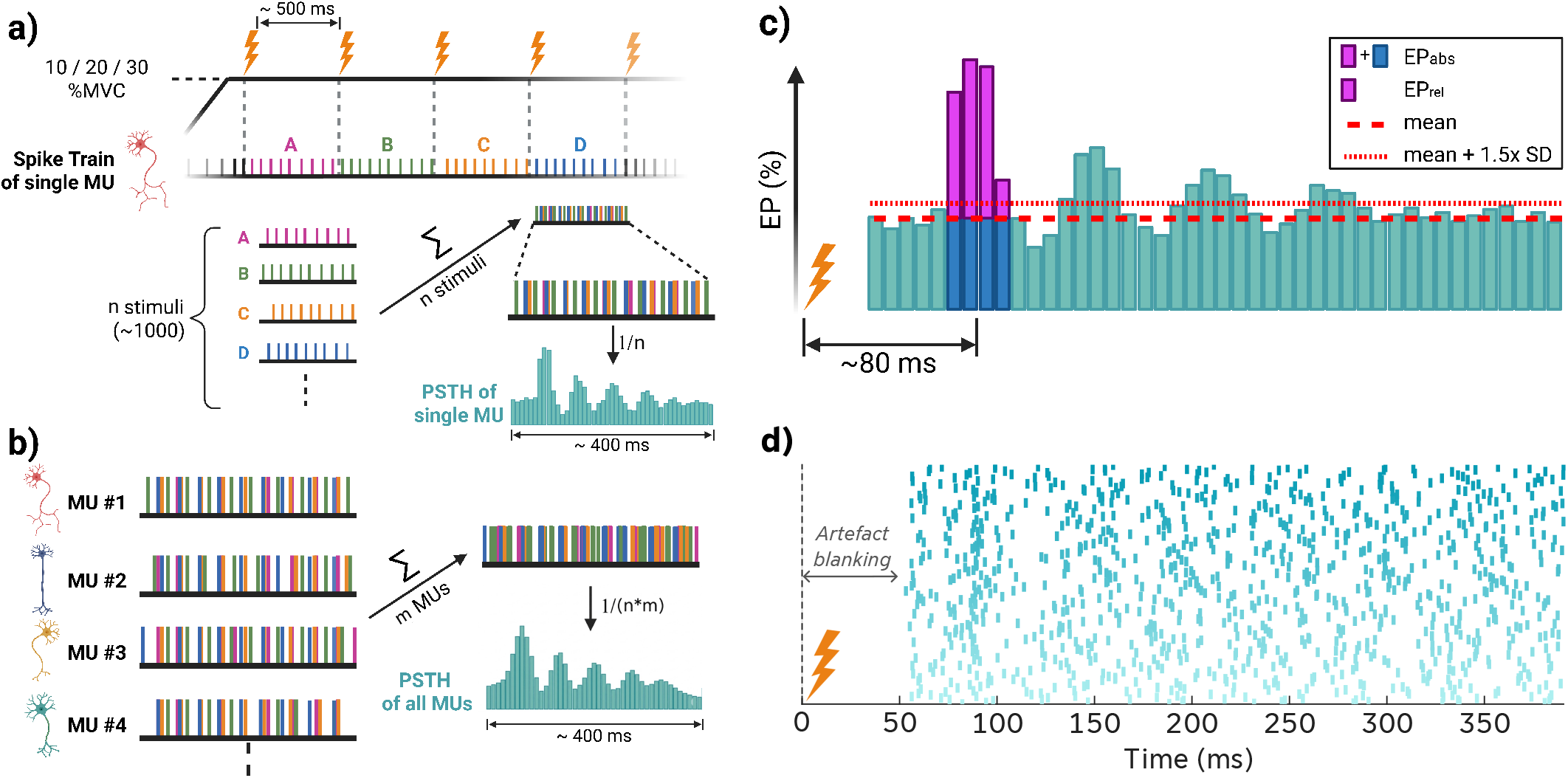
a) Computation of the PSTH from the spike trains of a single MU and the stimulation pulses. The PSTH is computed separately for each MU and the three force levels. b) Computation of the PSTH for the entire motor pool. c) Schematic illustration of a PSTH of MU firings displaying the reflex appearing due to cutaneous stimulation. The height of each bar represents the firing probability per MU within a given time range following a single stimulus. The sum of the purple bars indicates a reflex response, relative to the pre-stimulus mean (EP_*rel*_), typically occurring between 70 and 95 ms after stimulation [16]. The sum of the purple and dark blue bars displays the EP_*abs*_. The coarse dashed lines represent the pre-stimulus mean firing probability, while the fine dashed lines indicate 1.5 times the standard deviation above and below the mean. d) Exemplary raster plot of a single MU’s discharge pattern aligned to stimulus onset (0 ms). Data represent approximately 1000 stimulation trials. The period containing the stimulation artifact (0–50 ms) was blanked.

#### Population-level PSTH (Figure 3b)

In this approach, all MUs were considered jointly. For each stimulus, the relative discharge times of all identified MUs were computed with respect to the stimulus onset. The 400 ms post-stimulus window was divided into non-overlapping 5 ms bins. The number of discharges in each bin was counted across all MUs and all stimuli. The PSTH was then computed as:

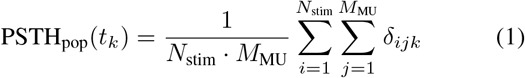

Here, *t*_*k*_ denotes the center of the *k*-th time bin, *N*_stim_ is the total number of stimuli, and *M*_MU_ is the total number of identified MUs. The binary indicator function *δ*_*ijk*_ equals 1 if MU *j* fired within bin *t*_*k*_ following stimulus *i*, and 0 otherwise.

#### Unit-specific PSTH (Figure 3a)

Alternatively, PSTHs were computed separately for each individual MU. This approach preserves unit-specific temporal discharge patterns. The PSTH for MU *j* was defined as:

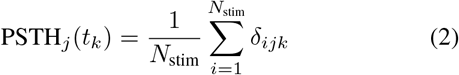

In this case, *δ*_*ijk*_ is again 1 if MU *j* fired in time bin *t*_*k*_ following stimulus *i*, and 0 otherwise. The result is a separate PSTH profile for each MU, normalized only by the number of stimuli.

To quantify the intensity of the evoked reflex response, the previously computed PSTHs were further analyzed. For each PSTH, all histogram bins within the expected reflex time window (60 - 120 ms post-stimulus) that exceeded a threshold defined as the mean plus 1.5 times the standard deviation were identified. The mean and standard deviation were calculated from a late, reflex-free segment of the PSTH. In previous studies, a threshold of two times the standard deviation has commonly been applied to identify the presence of a reflex response [35]. However, in the present study, prolonged PEW deflections following the reflex, manifested as delayed return of the PSTH to baseline, were occasionally observed. These extended deflections increased the variability in the later time window. To account for this and to maintain sensitivity in reflex detection, a more conservative threshold of 1.5 times the standard deviation was employed in this work. Based on the identified bins, two types of excitation probability (EP) were computed. The absolute excitation probability (EP_*abs*_) was defined as the sum of all bins exceeding the threshold, thereby reflecting the number of time intervals with elevated MU firing activity following stimulation. The relative excitation probability (EP_*rel*_) was calculated by subtracting the baseline mean from each of the identified bins prior to summation, thus providing a measure of the relative increase in firing probability above baseline activity.

These two EP metrics were derived separately for two types of PSTHs. For the population-level PSTH, which includes firings from all identified MUs, the resulting parameters were referred to as population-level absolute excitation probability (P-EP_*abs*_) and population-level relative excitation probability (P-EP_*rel*_). In contrast, for the unit-specific PSTH, where a separate PSTH was computed for each MU individually, the absolute and relative excitation probabilities were calculated for each MU separately and then averaged across all units. These unit-wise parameters were denoted as unit-level absolute excitation probability (U-EP_*abs*_) and unit-level relative excitation probability (U-EP_*rel*_). Consequently, four distinct reflex-related parameters were extracted and analysed per subject and force level: P-EP_*abs*_, P-EP_*rel*_, U-EP_*abs*_, and U-EP_*rel*_.

Subsequently, it was examined whether there is a correlation between EP and the recruitment threshold of individual motoneurons. For this analysis, U-EP_*rel*_ was used as the excitation probability measure, based on the assumption that U-EP_*abs*_ and U-EP_*rel*_ exhibit a similar trend in the preceding analysis (see Results section). An additional reason for using U-EP_*rel*_ is that it accounts for the influence of background firing of the MUs on the EP. The recruitment threshold for each MU was determined for each subject based on the most uniform ramp phase (out of the 10 ramps per force level). Specifically, the recruitment threshold was defined as the %MVC force at the time of the first firing of the respective motoneuron.

Furthermore, the change of U-EP_*abs*_ for the same MUs across different force levels was investigated. To this end, MUs were tracked across two recordings (from 10 to 20 %MVC and from 20 to 30 %MVC) based on their individual motor unit action potential (MUAP) shapes. For each MU, the MUAP shape was computed and compared to the shapes of MUs from the corresponding recording. The root mean square error (RMSE) between MUAP shapes was calculated to quantify similarity. This procedure, combined with careful manual inspection, allowed for the identification of MUs that were present in both recordings. Consequently, it was possible to assess how U-EP_*abs*_ of identical MUs changes with increasing force levels (Figure 4). The change in U-EP_*abs*_ of a MU across two force levels is referred to here as ΔU-EP_*abs*_.

**Fig. 4:**
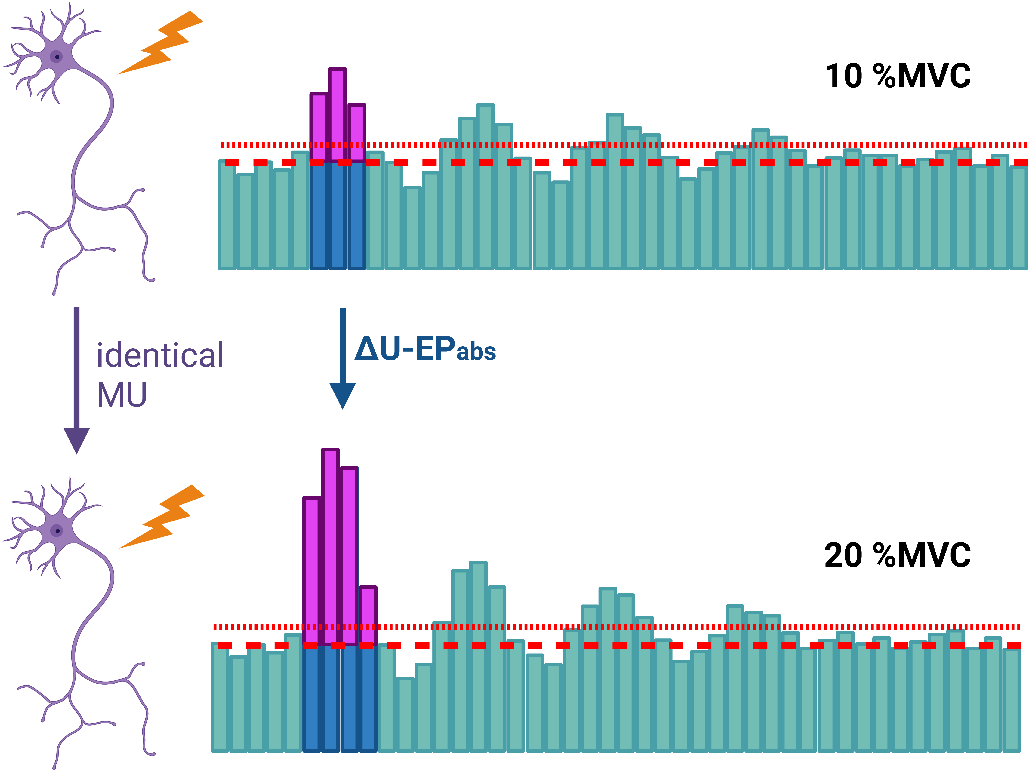
Schematic illustration of the change in U-EP_*abs*_ of a tracked MU at two different force levels. In this case ΔU-EP_*abs*_ is defined as the difference between U-EP_*abs*_ at 20 %MVC and U-EP_*abs*_ at 10 %MVC.

The typical shape of the PSTH related to the cutaneous reflex examined here includes a pronounced excitation occurring approximately 70–95 ms after the stimulus [16] (Figure 3c). This is typically followed by a decaying wave, here called PEW, with the most prominent negative deflection occurring immediately after the reflex-induced excitation. It is hypothesized that these PEWs arise from synchronization of MUs. Specifically, it is assumed when multiple MUs fire simultaneously as a result of the reflex, their firing patterns are reset and subsequently resume at the same firing frequency as before.

If this hypothesis holds true, there should be a correlation between the EP of individual MUs and the magnitude of the subsequent PEW. To investigate this, an individual PSTH is computed for each MU. To estimate the amplitude of the PEW, the first negative deflection, hereafter referred to as PED, is analyzed. For this purpose, the area between the pre-stimulus mean and the bars representing the firing probability is calculated for each MU. To account for inter-subject variability, the resulting PED magnitudes were Z-score normalized within each subject separately. This approach allows for examining whether a relationship exists between the EP of a single MU and the magnitude of the following negative deflection in its PSTH.

However, since correlation alone does not provide causal proof of synchronization-induced resetting, a targeted reflex-removal analysis was implemented. To validate this approach and establish a baseline for a pure synchronization-driven effect, a control dataset of 250 simulated MUs was generated. Each MU was modeled as a renewal process, where the ISIs followed a Gaussian distribution. The mean ISI (*µ*_*ISI*_) for each unit was derived from its firing rate (10–20 Hz), and the standard deviation was defined as *σ*_*ISI*_ = *µ*_*ISI*_ *CV* to impose a constant coefficient of variation (CV) of 0.12. To ensure physiological plausibility, a depression period was implemented by truncating the distribution at a minimum ISI of 20 ms. In accordance with the synchronization-resetting hypothesis, the reflex was simulated by inducing a phase reset: if a reflex-event (latencies: 70–95 ms, probability: 5– 30%) occurred before the next scheduled natural discharge, the existing ISI was truncated, a spike was registered at the reflex latency, and the timing of the subsequent spike was recalculated starting from this new reference point. Crucially, no independent inhibitory components were included in the simulation; any observed post-excitatory depression in the simulated PSTHs resulted solely from the reset-induced synchronization.

For both recorded and simulated MUs, an iterative greedy optimization algorithm was used to selectively discard stimuli associated with firing events within the reflex latency window. This process was repeated until the excitation peak in the resulting PSTH was reduced to the pre-stimulus baseline level. By comparing the residual magnitude of the PED in the “reflex-removed” PSTHs of simulated data (where the ground truth of pure synchronization is known) with that of the recorded data, it could be assessed whether the physiological negative deflection represents an independent inhibitory process or is a statistical consequence of the preceding excitation. To account for inter-subject variability in PED magnitudes while preserving the interpretability of the zero-line, the data were normalized using a median-scaling approach. For each subject, PED values were scaled relative to the median PED of the original (pre-removal) data. This normalization ensures that baseline differences between subjects are minimized without shifting the physiological meaning of the zero-reference, which represents a firing probability equal to the pre-stimulus mean.

## III. Results

In total, 854 MUs were identified from the right tibialis anterior of the nine subjects at the three contraction forces. This number includes potential common MUs that were identified at several contraction forces.

First, an analysis was performed to determine an appropriate number of stimuli to include. For this purpose, the U-EP_*rel*_ of each MU and the P-EP_*rel*_ of the entire motor pool were calculated separately for varying numbers of incorporated stimuli. To assess potential temporal effects, these calculations were performed using two distinct selection methods: sequential inclusion (chronological order) and random selection of stimuli. Figure 5 illustrates the average absolute change in U-EP_*rel*_ and P-EP_*rel*_ relative to the preceding value, expressed as a percentage of the final value, in increments of 50 stimuli. The colored lines (purple for U-EP_*rel*_, blue for P-EP_*rel*_) represent the sequential selection, while the gray lines depict the random selection.

**Fig. 5:**
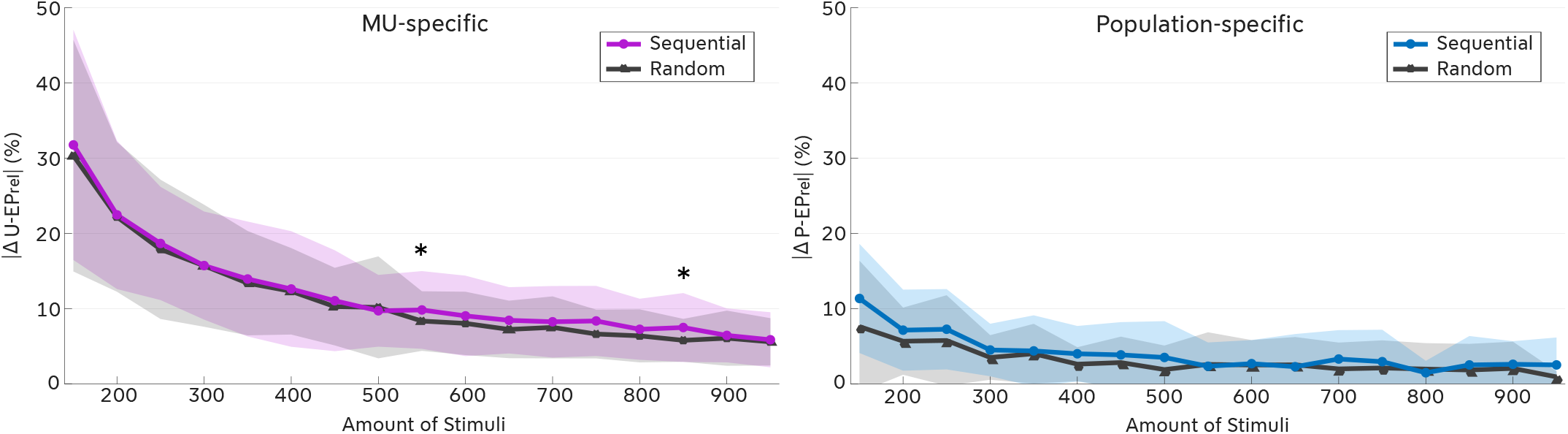
Average absolute change in excitation probability as a function of the number of included stimuli. The left panel illustrates the change in U-EP_*rel*_ for individual MUs, while the right panel displays the change in P-EP_*rel*_ for the aggregated motor pool. Purple (left) and blue (right) lines represent the sequential (chronological) selection of stimuli, whereas dark gray lines represent a random selection. Data are presented as the mean across all subjects, with shaded areas indicating the standard deviation (SD). The y-axis represents the absolute difference (Δ) relative to the preceding step (increments of 50 stimuli), expressed as a percentage of the final value. Asterisks (*) denote statistically significant differences between the sequential and random methods (*p* < 0.05, Wilcoxon signed-rank test with Bonferroni-Holm correction).

Both metrics exhibit a decaying trajectory, indicating that the variation between estimation steps decreases as the number of included stimuli increases. Notably, the magnitude of change is considerably larger for individual MUs compared to the aggregated motor pool. Statistical comparison between the sequential and random approaches was performed using a Wilcoxon signed-rank test with Bonferroni-Holm correction for multiple comparisons. Significant differences between the sequential and random approaches were observed only at two specific data points for the individual MUs: 550 (*p* = 0.024) and 850 stimuli (*p* = 0.034). No significant differences were found at any other stimulus count for single MUs, nor at any point for the motor pool. Within the range of 150–300 stimuli, which is commonly used in previous studies [1, 16, 21, 30, 31], the average change for individual MUs amounts to approximately 15–30%, whereas the motor pool shows a variation of approximately 5–10%. However, when substantially more stimuli are included, as in our dataset with up to 1000 stimuli, the average change is reduced to ∼5% for individual MUs and below ∼3% for the motor pool, indicating a smaller variation in excitation probability across different stimulus counts.

The analysis of reflex elicitation probabilities across the three force levels (10, 20, and 30 %MVC) revealed significant effects for all four PSTH-derived reflex measures. As shown in Figure 6a-d and Table 1, EPs increased systematically with higher force levels for all measures (P-EP_*abs*_, P-EP_*rel*_, U-EP_*abs*_, U-EP_*rel*_). Mean and standard deviation (*±* SD) values across subjects are reported in Table 1.

**TABLE I.**
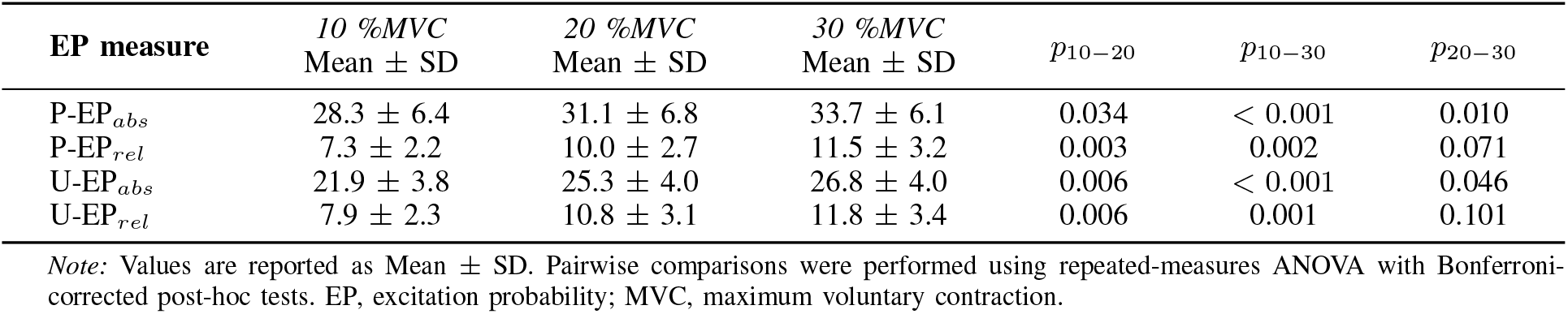
Excitation probabilities for three target force levels.

| EP measure | 10 %MVC<br>Mean ± SD | 20 %MVC<br>Mean ± SD | 30 %MVC<br>Mean ± SD | $p_{10-20}$ | $p_{10-30}$ | $p_{20-30}$ |
| --- | --- | --- | --- | --- | --- | --- |
| P-EP<sub>abs</sub> | 28.3 ± 6.4 | 31.1 ± 6.8 | 33.7 ± 6.1 | 0.034 | < 0.001 | 0.010 |
| P-EP<sub>rel</sub> | 7.3 ± 2.2 | 10.0 ± 2.7 | 11.5 ± 3.2 | 0.003 | 0.002 | 0.071 |
| U-EP<sub>abs</sub> | 21.9 ± 3.8 | 25.3 ± 4.0 | 26.8 ± 4.0 | 0.006 | < 0.001 | 0.046 |
| U-EP<sub>rel</sub> | 7.9 ± 2.3 | 10.8 ± 3.1 | 11.8 ± 3.4 | 0.006 | 0.001 | 0.101 |

Note: Values are reported as Mean ± SD. Pairwise comparisons were performed using repeated-measures ANOVA with Bonferroni-corrected post-hoc tests. EP, excitation probability; MVC, maximum voluntary contraction.

**Fig. 6:**
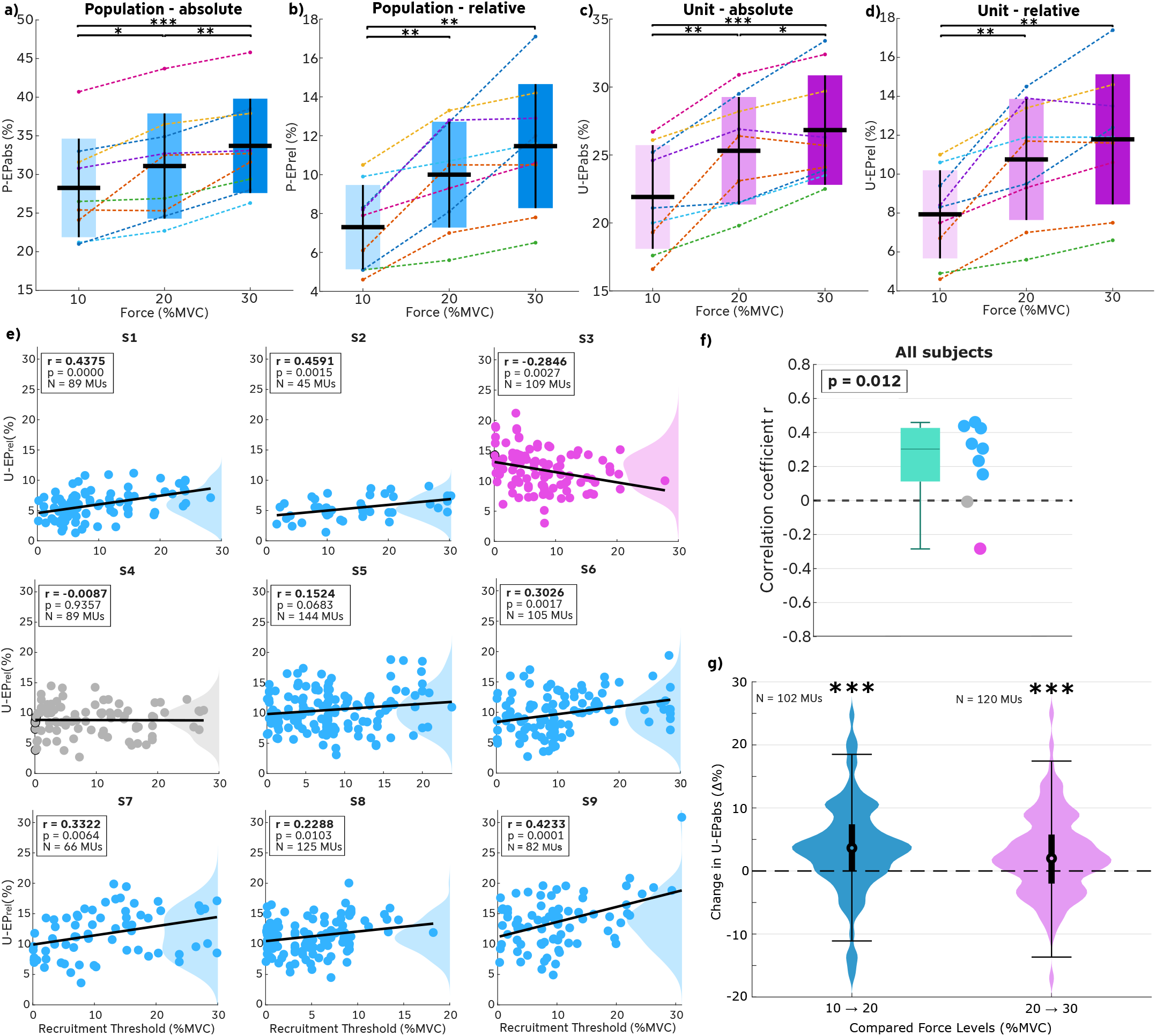
Post-stimulus excitation probability at different force levels calculated as a) P-EP_*abs*_ b) P-EP_*rel*_ c) U-EP_*abs*_ d) P-EP_*rel*_. Significance bars show the results of Bonferroni-corrected post-hoc tests. *** indicate *p* < 0.001, ** indicate *p* < 0.01, * indicate *p* < 0.05. e) Correlation between recruitment threshold and relative excitation probability for individual MUs, shown separately for each subject. Each dot represents one MU. The solid lines indicate the fitted linear regression lines. The shaded areas along the right y-axes represent the kernel density estimation (KDE) of the U-EP_*rel*_. f) Box plot summarizing the correlation coefficients across all subjects. Blue dots indicate positive correlation coefficients, grey dots indicate coefficients near zero, and pink dots indicate negative correlation coefficients. g) Change in U-EP_*abs*_ of tracked MUs (ΔU-EP_*abs*_) across different force levels. Shown are the changes between 10 and 20 %MVC as well as between 20 and 30 %MVC. The violin shape represents the probability density function (distribution) of the data. The distribution is summarized by an embedded box plot: The thick vertical line shows the Interquartile Range (IQR), spanning from the 25^th^ (*Q*_1_) to the 75^th^ (*Q*_3_) percentile. The central white marker indicates the Median. The thin vertical lines (whiskers) extend to the most extreme data points not considered outliers (typically 1.5 × IQR). Statistical significance against zero (*µ* = 0) was assessed using a one-sided T-test (testing the hypothesis *µ* > 0). Asterisks (***) indicate statistical significance at *p* < 0.001 for both groups. The comparison between the 10 → 20 %*MV C* and 20 → 30 %*MV C* groups was assessed using the Mann-Whitney U-Test and showed no significant difference (*p* = 0.093).

Pairwise comparisons confirmed significant differences between 10 %MVC and both 20 %MVC and 30 %MVC for all four measures (all *P ≤* 0.034). Regarding the transition from 20 %MVC to 30 %MVC, significant increases were observed for the absolute elicitation probabilities (P-EP_*abs*_, *P* = 0.010; U-EP_*abs*_, *P* = 0.046). In contrast, the relative measures did not reach statistical significance for this comparison (P-EP_*rel*_, *P* = 0.071; U-EP_*rel*_, *P* = 0.101), although a trend toward higher values was visible.

Statistical differences between force levels were tested using a repeated-measures ANOVA. Sphericity was assessed using Mauchly’s test, and the assumption was met for all measures. Post-hoc pairwise comparisons were performed with Bonferroni correction.

The scatter plots (Figure 6e) illustrate, for each participant separately, the relationship between U-EP_*rel*_ and the recruitment threshold of individual MUs. Each dot represents a single MU. Linear regression lines were fitted to the data to visualize the slope of the correlation between the two parameters. Statistical significance was assessed using Pearson’s correlation. The results show that 7 out of 9 participants (∼78 %) exhibited a positive correlation between recruitment threshold and U-EP_*rel*_ of individual MUs (0.15 < *r* < 0.5). Except for one participant (S5), these correlations were statistically significant (*p* < 0.05). One participant (S4) showed no correlation at all (*r* = −0.009, *p* = 0.936), while another (S3) exhibited a significant negative correlation (*r* = −0.2846, *p* < 0.01).

The summary of all correlation coefficients is shown in the box plot (Figure 6f). A one-sided (right-tailed) t-test was used to determine whether the correlation coefficients were significantly greater than zero. The results indicate that the coefficients were indeed significantly greater than zero (*p* < 0.05).

A total of 102 unique MUs were successfully tracked between the 10 and 20 % MVC recordings across all subjects. For the transition between 20 and 30 % MVC, 120 identical MUs were detected. The change in U-EP_*abs*_ was calculated for each matched MU pair between the respective contraction levels (Figure 6g). The median change (ΔU-EP_*abs*_) from 10 to 20 % MVC was 3.65% (IQR: 0.00% to 7.40%), which was statistically highly significant (*p* = 7.69 × 10^−7^). Similarly, the median change from 20 to 30 % MVC was 2.00% (IQR: − 2.00% to 5.78%), also reaching a high level of statistical significance (*p* = 2.38 × 10^−5^). The difference in median change between the 10 → 20 %*MV C* and 20 → 30 %*MV C* transitions was not statistically significant (Mann-Whitney U-Test, *p* = 0.093). The *p*-values were obtained using one-sided, one-sample T-tests, testing the hypothesis that the mean change (*µ*) in ΔU-EP_*abs*_ of the tracked MUs between force levels was significantly greater than zero (*µ* > 0).

The scatter plots shown in Figure 7a illustrate the relationship between the z-score normalized magnitude of the PED and EP for individual MUs. The left plot of Figure 7a uses the U-EP_*abs*_ while the right one shows the correlation to the U-EP_*rel*_. Each data point represents a single MU, and each color corresponds to all MUs from one subject. A linear regression line was fitted to the data, revealing a positive association between the two variables for both, the absolute and relative EPs. PED magnitude correlated moderately and significantly with both U-EP_*abs*_ (*r* = 0.4102, *p* < 0.001) and U-EP_*rel*_ (*r* = 0.5041, *p* < 0.001). In both cases, the Pearson correlation coefficients indicated a robust positive relationship.

**Fig. 7:**
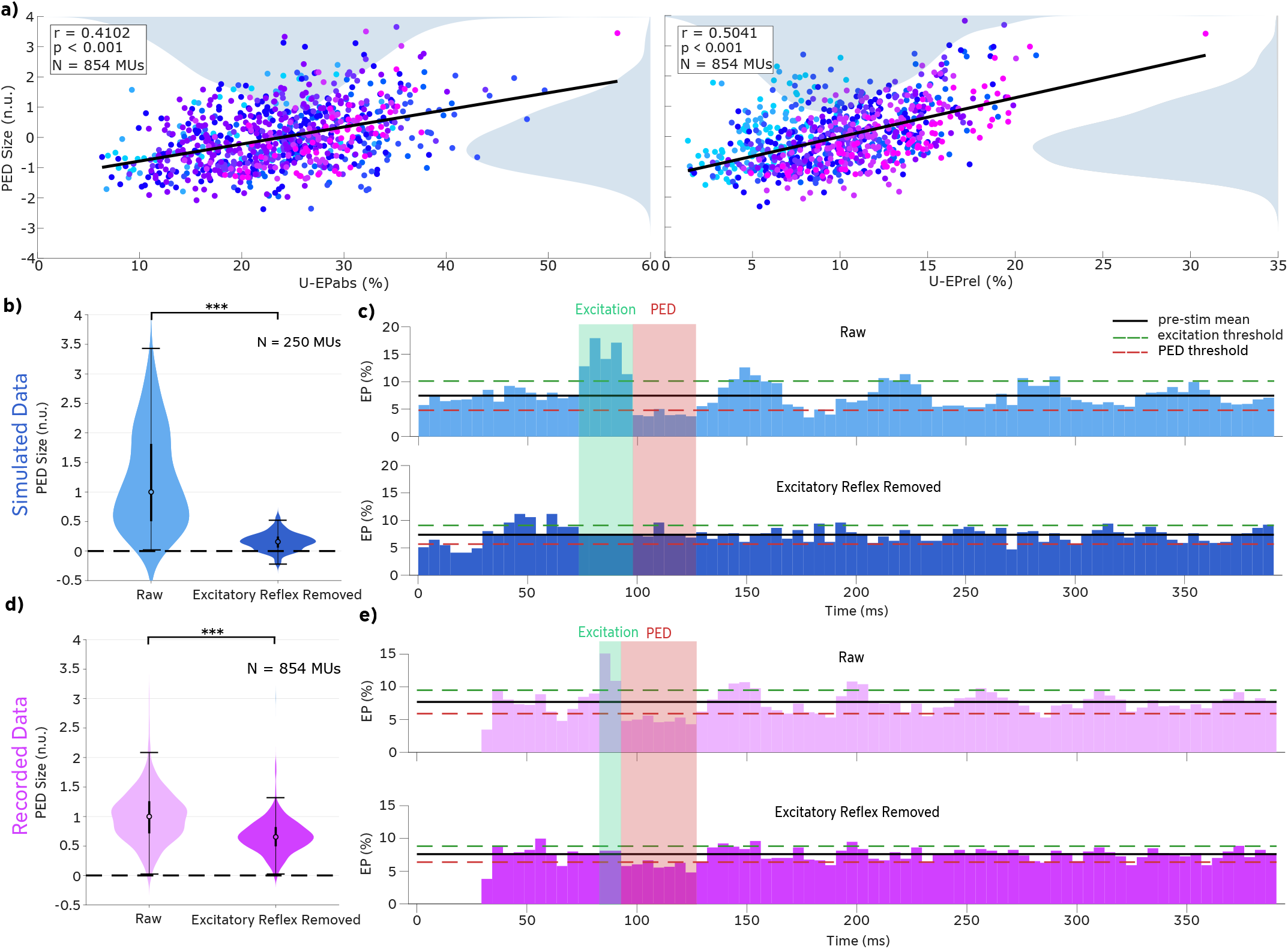
Post-excitation depression (PED) size (z-normalized) versus absolute (top) and relative (bottom) excitation probability of individual MUs. Each color represents all identified MUs pooled from a single subject. The solid line indicates the fitted linear regression line, showing a positive slope. Marginal KDE plots show the distribution of data along the x- and y-axes. The reported *p*-values (*p* < 0.001) were calculated using Pearson’s correlation coefficient (*r*) and confirms a highly significant positive correlation between the post-excitation depression period magnitude and the excitation probability of individual MUs. b) Violin plots showing the distribution of PED sizes for 250 simulated MUs in the raw state compared to the state after targeted excitatory reflex removal. Values are normalized using a median-scaling approach to preserve the physiological reference of the baseline. Statistical significance between conditions was assessed using a Wilcoxon signed-rank test (*** indicate *p* < 0.001) c) Exemplary PSTH of a single simulated MU before (top) and after (bottom) excitatory reflex removal. Note that after removal, neither a discernible relative excitation nor a subsequent PED is present. d) Distribution of PED sizes for recorded data (854 MUs) displayed as violin plots, following the same methodology and median-scaling as in panel b) (Wilcoxon signed-rank test; *** indicate *p* < 0.001). e) Exemplary PSTH of a recorded MU before (top) and after (bottom) targeted excitatory reflex removal.

**Fig. 8:**
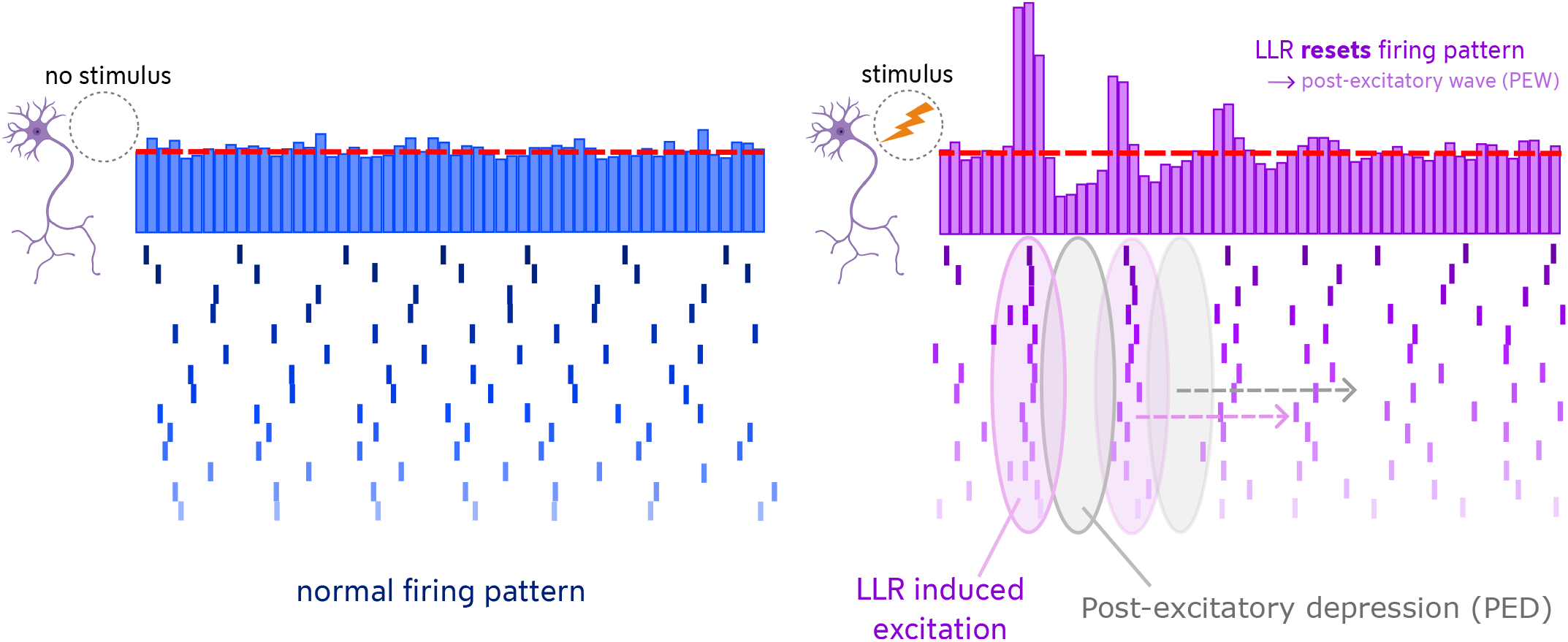
Modulation of concurrent MU firing patterns by sensory stimulation. The panels illustrate MU discharge patterns without stimulation (left) and with sensory stimulation (right). The excitation evoked by the LLR synchronizes the firing times of the MUs, resulting in a post-excitatory depression (PED) affected by this synchronization visible in the PSTH. This observation indicates that the LLR resets the intrinsic firing cycle of the MUs rather than merely inserting an additional spike into the ongoing train. Consequently, a decaying oscillatory post-excitatory wave (PEW) is observed in the PSTH. The red dashed lines indicate the pre-stimulus means.

To investigate whether these observed depressions are a consequence of firing-pattern resetting or represent an independent inhibitory process, a targeted excitatory reflex removal analysis was performed (Figure 7b–e). In the simulated dataset (250 MUs), where the PED was exclusively generated by synchronization-induced resetting, the removal of reflex-associated stimuli led to a near-complete elimination of the negative deflection. The median PED magnitude decreased by 84.2%, dropping from 1.000 [IQR: 0.505, 1.815] to 0.158 [IQR: 0.059, 0.246] (Wilcoxon signed-rank test, *p* < 0.001). This effect is clearly visible in the exemplary PSTH (Figure 7c), where the post-reflex firing probability returns to baseline levels once the synchronization peak is removed.

In contrast, the reduction observed in the recorded MU data (854 MUs) was considerably less pronounced. Although the reduction was statistically highly significant (Wilcoxon signed-rank test, *p* < 0.001), the median magnitude only decreased by 34.7%, from 1.000 [IQR: 0.713, 1.260] to 0.653 [IQR: 0.494, 0.821]. This discrepancy is further illustrated by the exemplary PSTH in Figure 7e: unlike the simulated data, the recorded MU still exhibits a distinct and persistent dip at the PED latency even after the reflex-induced spikes have been eliminated.

## IV. Discussion

To investigate our hypothesis that LLR responses are shaped by the combined integration of spinal motor neuron excitability and the baseline net neural drive, this study provided a high-resolution characterization of cutaneously-elicited LLRs at the level of single MUs. Our findings demonstrate that MU reflex responses are not uniform but exhibit significant heterogeneity depending on MU-specific characteristics. We analyzed the modulation of reflex excitability across different levels of voluntary contraction. We observed a systematic increase in reflex elicitation probabilities with increasing force levels (10% to 30% MVC). Given the lack of a standardized consensus in the literature on how to best quantify reflex magnitude, whether calculated as an aggregate of the motor pool (normalized to the number of identified units) or averaged across single unit estimates, and whether expressed as absolute values or relative to background activity, we employed a multi-faceted approach consisting of four distinct measures (P-EP_*abs*_, P-EP_*rel*_, U-EP_*abs*_, U-EP_*rel*_).

The fact that all four metrics yielded a consistent positive correlation with contraction level underscores the robustness of this phenomenon. Regardless of the calculation method, the LLR response intensifies as the voluntary neural drive to the muscle increases. This finding aligns with previous observations describing a gain scaling of both cutaneously and Ia-afferent mediated reflex responses with voluntary contraction force [35, 36]. This consistency serves as a validation of our experimental setup, confirming that our data reproduces established physiological trends before delving into the more granular MU-specific mechanisms.

It is, however, noteworthy that while absolute measures continued to increase significantly between 20 and 30 %MVC, the relative measures (normalized to background activity) did not exhibit significant further growth in this range. This suggests that while the absolute reflex output per MU grows with force, likely due to higher firing rates, it scales proportionally with the concurrent background neural drive at higher force levels, resulting in a stable relative magnitude. A similar dependency of the reflex magnitude, the so-called automatic gain-scaling, on the applied voluntary background force has been extensively documented for spinal SLRs, as in stretch reflexes, in previous studies [37–41].

To understand the physiological mechanisms underlying the observed gain scaling of reflex responses with contraction force, we investigated two potential contributing factors: the specific recruitment of MUs with higher reflex susceptibility, and the intrinsic modulation of already active units.

First, our analysis of the relationship between recruitment threshold and reflex excitability revealed a significant positive correlation across the subject group (*p* = 0.012). Contrary to our initial hypothesis that LLR modulation would follow a continuum consistent with Henneman’s size principle, where smaller, low-threshold motor neurons typically exhibit higher synaptic gain, our results demonstrate an inverted relationship. In the majority of participants (∼78 %), MUs with higher recruitment thresholds exhibited a higher probability of discharging a reflex spike.

This finding stands in contrast to observations in the upper limb, where Garnett and Stephens [42] reported in 1980 that high-threshold units in the first dorsal interosseous muscle exhibited lower excitation probabilities following digital nerve stimulation. Notably, only one of our nine participants significantly followed the pattern observed by Garnett and Stephens, whereas the majority showed the opposite trend. The exact physiological basis for this discrepancy remains to be elucidated; it may stem from intrinsic differences in the synaptic organization between the intrinsic hand muscles and the lower limb dorsiflexors, or methodological variations in stimulation. Regardless of the underlying mechanism, our results indicate that for the tibialis anterior, the central nervous system does not engage the motor pool uniformly during a reflex response. Instead, high-threshold MUs, which are typically recruited later according to Henneman’s size principle [43, 44], appear to possess a greater responsiveness to the cutaneous synaptic input in the lower limb. Consequently, as voluntary force increases and these high-threshold units are recruited into the active pool, they disproportionately contribute to the overall increase in the reflex magnitude observed at the population level.

However, the moderate strength of the positive correlations (*r* ranging from 0.15 to 0.45) and the inter-subject variability, including one subject with no correlation and one with a significant negative relationship, indicate that the recruitment threshold is a significant predictor but not the sole determinant of reflex excitability. However, it is important to note that our voluntary force level was up to 30 %MVC, while the tibialis anterior muscle has a full MU recruitment range of up to nearly 90 %MVC [45]. Therefore, our findings might be limited to the behavior of lower-threshold MUs and may not necessarily be generalizable to the muscle’s full functional breadth. This heterogeneity implies that the synaptic weighting of cutaneous afferents onto the motor pool is not strictly determined by the size principle alone. Indeed, deviations from the standard voluntary recruitment order are not unprecedented. Studies comparing voluntary versus electrically evoked contractions have demonstrated that the recruitment order can be random or even reversed in a substantial proportion (28–35%) of MU pairs [46]. This supports the perspective that the MU activation hierarchy is flexible and contingent upon the specific source of neural drive. Therefore, other factors, such as the specific topological distribution of afferent terminals or intrinsic differences in bistable firing behaviors, likely contribute to the variance in excitability that is not explained by the recruitment threshold alone.

Second, beyond the recruitment of new units, we isolated the effect of increased neural drive on individual MUs by tracking the same units across force levels. This analysis demonstrated that identical MUs significantly increase their excitation probability when the background contraction, and thus the net synaptic drive, increases. This finding is crucial as it decouples the unit’s “identity” from its “state.” It shows that the increase in reflex magnitude is not solely due to the recruitment of “more sensitive” units (as discussed above), but also because the elevated net excitatory drive brings the membrane potential of active MUs closer to their discharge threshold. In this state, the same postsynaptic potential elicited by the cutaneous stimulus is more likely to cross the threshold and trigger an action potential.

In summary, the systematic increase of reflex excitability with voluntary force is driven by the interplay of at least two identified mechanisms: (1) the progressive recruitment of high-threshold MUs that are inherently more excitable in response to this specific stimulus, and (2) the increased discharge probability of the active motor pool due to a heightened background of excitatory drive. However, given the remaining variance in our data, it is highly probable that additional, yet to be identified factors, such as non-uniform synaptic weighting or intrinsic motoneuron properties, further contribute to the modulation of MU excitability. Moreover, it may be hypothesised that the underlying MU modes may also change excitability of specific MUs [47].

Finally, we addressed the physiological origin of the PEW typically observed in the PSTH and EMG signals following the LLR. Driven by the observed post-excitatory discharge patterns, our third, a posteriori hypothesis posited that the signal decline immediately succeeding the reflex peak is primarily a deterministic consequence of the discharge reset rather than a distinct, centrally mediated inhibitory component. Our analysis revealed a highly significant positive correlation (*p* < 0.001) for both cases: both absolute and relative EP of individual motor neurons correlated with the magnitude of the subsequent synchronization-induced PED. This finding indicates that MUs with a higher likelihood of discharging an LLR spike exhibit a proportionally stronger “silence” immediately thereafter. This relationship supports the “synchronization-reset” mechanism: when a cutaneous stimulus synchronizes a fraction of the motor pool to fire within a narrow time window (the LLR peak), these units simultaneously enter their after-hyperpolarization phase (Figure 8).

To transition from correlation to causal evidence, we implemented a novel reflex-removal analysis. The validity of this method was first established using a simulated dataset (250 MUs), where the PED and PEW were exclusively gen-erated by firing-pattern resetting. In these simulated units, the targeted removal of reflex-associated spikes eliminated the subsequent depression by 84.2%, effectively returning the PSTH to baseline levels (Figure 7c). This near-complete elimination confirms that the algorithm successfully isolates and removes the statistical consequences of synchronization.

However, applying this same procedure to the recorded physiological data (854 MUs) yielded a more nuanced result. While we observed a highly significant reduction in PED magnitude (*p* < 0.001), the reduction was considerably less pronounced than in the simulation, amounting to only 34.7% (Figure 7d). As illustrated in the exemplary recorded PSTH (Figure 7e), a distinct and persistent dip remains even after the excitatory reflex component is removed. This discrepancy suggests that while synchronization-resetting is a substantial contributor to the observed depression, it does not fully account for the PED in real motor unit data.

Consequently, the post-reflex “inhibition” should be viewed as a hybrid phenomenon. While a significant portion is indeed an intrinsic artifact of the synchronized resetting of the firing cycle, the persistent deflection observed in recorded units indicates the likely superimposition of a genuine, centrally mediated inhibitory synaptic input. This distinction is crucial for interpreting LLR PEWs, as it implies that the declining phase reflects both a recovery process and a separate modulation of neural drive. Crucially, the subsequent return to baseline manifests as a decaying oscillation. This damping occurs because the synchronized MUs possess heterogeneous discharge rates [48], causing their spike timing to drift apart over subsequent cycles (temporal dispersion). Furthermore, the inherent physiological variability in the discharge frequency of individual units [49] contributes to this desynchronization, explaining why the oscillatory pattern fades over time even in single-unit PSTHs.

Lastly, for the methodological validation of our stimulus protocol, our data reveals that accurately capturing these subtle physiological differences requires a methodological approach that goes beyond current standards. We show that the stability of reflex parameter estimation is heavily dependent on the number of stimuli applied, especially to single MUs. Consequently, the physiological insights presented in this work are inextricably linked to a high-repetition protocol that ensures the separation of true physiological signal from stochastic noise, a factor that has likely been underestimated in previous literature.

A primary methodological objective of this study was to challenge the reliability of stimulus counts typically employed in single-MU reflex analysis. Previous studies have commonly relied on approximately 150 to 300 stimuli to estimate reflex parameters [1, 16, 21, 30, 31]. Our stability analysis demonstrates that within this range, the estimation of the excitation probability for single MUs (U-EP_*rel*_) remains volatile, exhibiting fluctuations of 15–30% with the addition of 50 further stimuli. In contrast, extending the protocol to approximately 1000 stimuli reduced this variation to ∼ 5%, indicating that a stable plateau for reliable estimation is reached only at significantly higher repetitions than previously assumed.

It is worth noting that this instability is less pronounced at the aggregate level. When analyzing the motor pool parameters (P-EP_*rel*_), the variation within the standard 150–300 stimulus range was considerably lower (approximately 5–10%) than for individual MUs. While the motor pool estimates also exhibited further stabilization with increased stimulus counts (dropping to < 3% variation), the contrast emphasizes that the sufficiency of the stimulus count depends on the resolution of the analysis. While lower counts might be debatable for coarse, aggregate motor pool assessments, our data strongly suggests they are inadequate for the high-granularity analysis of single MUs, where the stochastic variability is not smoothed out by population averaging.

This observation is critical because reflex parameters such as the EP are stochastic in nature rather than deterministic fix-values. Therefore, according to the law of large numbers, a larger sample size is mandatory to reduce the standard error of the probability estimate. The fact that standard protocols operate in a range where the single-MU estimator still fluctuates by up to 30% suggests that some variability attributed to physiological differences in prior studies might instead reflect methodological noise.

Importantly, our comparison of sequential versus random stimulus selection revealed no systematic differences across the vast majority of the protocol. This similarity indicates that the observed stabilization at higher stimulus counts is primarily driven by statistical convergence rather than temporal physiological effects such as habituation or fatigue. If fatigue were the dominant factor, the sequential analysis would have diverged significantly from the random selection. Thus, we conclude that the high-repetition approach employed here provides a methodological benchmark necessary for the robust interpretation of MU-specific reflex physiology.

While this study provides a robust methodological framework for analyzing MU-specific reflexes, certain limitations should be acknowledged. First, our investigation was restricted to the tibialis anterior muscle using electrical cutaneous stimulation. While this setup is a standard model for studying spinal reflexes, the generalizability of our findings to other muscle groups (e.g., upper limb muscles) or mechanical stimuli (e.g., stumbling perturbations) remains to be verified. Different neural circuits or afferent populations might exhibit distinct integration properties.

Second, the high-repetition protocol (*N* ≈ 1000), which we identified as necessary for reliable parameter estimation, imposes a significant time burden and requires sustained attention from participants to maintain steady isometric contractions. While we successfully controlled for fatigue and attention-related drift in our cohort, the practical feasibility of such extended protocols in clinical populations with motor impairments may be limited. Future research should explore adaptive estimation techniques or advanced signal processing methods that might achieve similar reliability with fewer stimuli.

## V. Conclusion

This study advances the understanding of sensorimotor integration by providing a high-resolution characterization of cutaneously-elicited LLRs at the single MU level. Physiologically, our data reveals that the gain scaling of the cutaneous reflex with voluntary force is driven by the interplay of at least two mechanisms: the recruitment of high-threshold MUs and the increased discharge probability of the active motor pool due to elevated net synaptic drive.

Furthermore, our results provide a more nuanced perspective on the post-excitatory depression in the discharge pattern.

By implementing a targeted reflex-removal analysis validated through motor unit simulations, we demonstrated that this suppression is a hybrid phenomenon. While a substantial portion of the observed depression is a deterministic consequence of the synchronized discharge reset, accounting for the majority of the effect in simulated data, the persistence of a significant negative deflection in recorded MUs suggests the additional involvement of an independent, centrally mediated inhibitory reflex process.

Methodologically, we demonstrated that the stochastic nature of reflex generation benefits significantly from higher stimulus counts than typically employed in the literature. Standard protocols using 150–300 repetitions were shown to yield estimates with higher variability for individual MUs, whereas increasing the count to *∼* 1000 substantially improved stability, yielding more precise and robust estimates. Collectively, these findings establish a new methodological benchmark for future reflex studies and offer deeper insights into the complex, multilayered control mechanisms of the human motor system.

## Additional Information

### Author contributions

Conceptualization: Y.F. and A.D.V. Data acquisition and curation: Y.F. Formal analysis: Y.F. Funding acquisition: A.D.V. Investigation: Y.F. Methodology: Y.F., D.S.S. and A.D.V. Project administration: Y.F. and A.D.V. Resources: Y.F. and A.D.V. Supervision: A.D.V. Writing-original draft: Y.F. Writing-review and editing: Y.F., D.S.S. and A.D.V.All authors have read and approved the final version of the manuscript.

